# Human decompression in real time: programmable ultrasound imaging during hyperbaric exposure

**DOI:** 10.64898/2026.07.22.737513

**Authors:** Joshua B. Currens, Michael J. Natoli, Katherine Eltz, Gabriela Morales, Kathlyne Jayne B. Bautista, Paul A. Dayton, Rachel M. Lance, Omer Oralkan, Feysel Yalcin Yamaner, Richard E. Moon, Virginie Papadopoulou

## Abstract

The formation of inert gas bubbles during decompression can lead to decompression sickness (DCS), a major operational risk for divers, compressed-gas workers, astronauts, and high-altitude aviators. In diving, DCS risk is typically inferred from post-dive ultrasound detection of venous gas emboli (VGE), precluding modification of decompression schedules based on real-time physiological feedback. Two-dimensional ultrasound imaging could provide additional insight into decompression-related physiological changes; however, its use in hyperbaric environments has been largely precluded by fire risk associated with elevated oxygen partial pressures (ppO₂) in enclosed spaces. Here, we developed a workflow for operating a programmable ultrasound system under hyperbaric conditions and acquiring ultrasound data from the subclavian vein and calf muscle during decompression. A total of 42 dives were conducted by 26 individuals using a previously characterized dive profile to 132 feet seawater (FSW) for 20 min with 9 min of decompression. Three exposure conditions were evaluated: non-exercising, exercising, and a brief pause at 20 FSW during compression. Twelve dives included programmable ultrasound imaging during decompression. Post-dive VGE responses were consistent with prior reports while demonstrating substantial inter-individual variability and sensitivity to modest profile modifications. VGE were detected in the subclavian vein during decompression in two participants and subsequently confirmed by post-dive echocardiography. Calf muscle ultrasound brightness typically increased from pre-dive to decompression measurements, before decreasing below baseline in the 120 min post dive measurement period. These findings demonstrate the feasibility of programmable ultrasound imaging during human decompression and establish a practical framework for ultrasound operation under hyperbaric conditions. This approach may support future physiological studies and development of automated decompression monitoring technologies.

**New and Noteworthy:** This study demonstrates the first use of a programmable ultrasound system to acquire and quantitatively analyze ultrasound data during human decompression. The approach enabled direct visualization of venous gas emboli during decompression and revealed calf muscle ultrasound signal changes, providing a new tool for investigating physiological responses during decompression that are not accessible through conventional post-dive monitoring.

## Introduction

Decompression sickness (DCS) is an ailment that can occur after a reduction in ambient pressure, examples of which include returning from an undersea dive, tunnel work, or preparing for extravehicular activity in space. The decrease in ambient pressure can cause dissolved inert gas within tissues and blood to come out of solution and form bubbles which, in excess, may cause DCS (1, 2). Those suffering from DCS present with various manifestations ranging in severity from mild skin rashes to paralysis, loss of consciousness and death. The standard treatment for DCS is prompt recompression in a hyperbaric chamber with pure oxygen breathing (2). DCS occurs in recreational scuba diving [∼1 in 10,000 dives, (3), depending on the circumstances], and one astronaut has reported knee pain consistent with DCS during depressurization in space (1, 4, 5). A higher probability of DCS is tolerated in US Navy dives of up to 2% for mild DCS (6), presenting an operational hazard. The risk of severe injury in a remote or austere location can jeopardize personnel, resources, and the operational objective. Current decompression schedules are derived from theoretical models, which are rigidly obtained for a given depth and time without accounting for other factors. Furthermore, only a limited amount of physiological monitoring and data has been collected during hyperbaric exposure. Conversely, wearable devices have grown in popularity and could provide a rich source of information for validating decompression procedures. Real-time diver monitoring during decompression could be incorporated into existing theoretical models to tailor the return to surface individually.

DCS risk mitigation is commonly achieved by a decompression algorithm that schedules the return to surface based on depth and time of the dive exposure. There are two categories of decompression algorithms, probabilistic and deterministic, both of which assume the same level of DCS risk for all divers using a population approach and do not incorporate real-time physiological status (7–9). As a result, the current framework for decompression planning is unable to account for the individual factors that may impact a diver’s susceptibility on a given day or dive exposure, which has been shown to vary (10–15). The population approach to dive planning may cause the assigned decompression obligation to be over- or under-prescribed in practice for the individual, resulting in DCS cases for dives that adhered to the models (3, 10). Efficient, personalized decompression strategies may ultimately be enabled by wearable physiological sensors, including emerging ultrasound imaging technologies capable of real-time monitoring during decompression.

Circulating bubbles, or venous gas emboli (VGE), detected by ultrasound are regularly used as an early study endpoint to estimate population DCS risk for a given dive profile While the absence of VGE is strongly associated with low DCS risk, elevated VGE grades have limited positive predictive value for individual DCS outcomes (10, 16). VGE are detected with ultrasound using either Doppler audio or 2-D echocardiography to identify the bubbles as chirps or localized hyperechoic regions, respectively. Research consensus guidelines require VGE to be measured at 20-minute intervals for at least two hours post-dive to characterize bubble presence at a given time, starting within 20 min of surfacing (17). Higher peak post-dive VGE presence has shown an increased DCS risk, suggesting that divers with more bubbles may be at a slightly higher risk of injury (16, 18), although the predictive power is relatively weak (19). To date, ultrasound data on divers is almost entirely limited to intravascular bubbles post-hyperbaric exposure, preventing researchers from studying physiological changes during the crucial decompression phase. A limited amount of hyperbaric chamber dives were previously conducted that included Doppler ultrasound monitoring during the decompression phase, which demonstrated VGE presence could occur prior to surfacing and would increase during as the decompression period progressed (20–23). Doolette & Mitchell have presented a review of Doppler measurements collected at pressure during experimental saturation dives in a hyperbaric chamber (24); however, most measurements were collected once the diver returned to storage pressure from an excursion, not during the decompression. The occurrence of VGE during decompression, which is not regularly monitored, demonstrates that post-dive VGE presence provides an incomplete picture. Previous hyperbaric decompression experimentation was limited to aurally monitoring the vasculature for VGE, which may or may not precede DCS symptoms (20). In hypobaric decompression exposures, VGE presence has been shown with Doppler (25) and two-dimensional echocardiography (4, 26). However, VGE detection with two-dimensional (2D) echocardiography requires extensive training and an ideal environment to collect reliable data, making it a suboptimal location for a wearable device to monitor decompression status. Additionally, the peak post-dive VGE presence has presented the strongest correlation at 10% for Doppler (18) and 5% for 2-D echocardiography (16). Identification of the peak EB grade requires the decompression to have already been completed, at which point schedule adjustments can no longer be made. Beyond VGE detection, investigations into quantitative ultrasound signal analysis have demonstrated early utility as a supplement to traditional B-mode imaging techniques in human (27) and animal models (28–30), detecting bubbles in soft tissue. Nonlinear ultrasound modes, including pulse inversion methods developed for contrast-enhanced vascular imaging, may improve contrast-to-tissue sensitivity and bubble detection (31). However, implementation of nonlinear ultrasound approaches in decompression research remains limited (30, 31).

To better characterize the physiological responses to decompression, we developed a workflow to acquire ultrasound measurements on human participants before, during, and after a simulated dive in a hyperbaric chamber with a research programmable ultrasound system to enable collection of pre-processed ultrasound data. A longstanding challenge in decompression research is the limited comparability between studies due to differences in dive profiles, environmental conditions, and other factors that influence post-dive outcomes. To facilitate comparison with prior work, we selected a dive exposure that has been extensively characterized by the U.S. Navy Experimental Diving Unit and is associated with a low risk of DCS while consistently producing substantial post-dive VGE (32). We therefore sought to evaluate the feasibility of programmable ultrasound imaging during decompression, characterize post-dive VGE responses to a well-established dive profile, determine whether VGE could be directly visualized during decompression using two-dimensional ultrasound imaging, and explore whether localized tissue ultrasound measurements exhibit detectable changes during decompression.

## Methods

### Ethical approval

All study procedures were conducted under protocols approved by the institutional review boards (IRB) of Duke University and of the University of North Carolina at Chapel Hill (IRB #Pro00111310 and #22-2449). All participants provided written informed consent prior to enrollment and participation in experiments. All procedures were performed in accordance with institutional guidelines and the Declaration of Helsinki.

### Participants and study design

Each participant underwent a medical screening and was cleared for hyperbaric exposure before enrollment. A total of 26 participants (20 men and 6 women) were included. Participant average age was 34 (range: 19 to 61) years and included a range of diving experience levels including no prior dive experience. Participants were instructed to refrain from diving for at least 24 hours prior to the experimental exposure and to report any post-exposure symptoms to the attending physician for medical evaluation and possible diagnosis of DCS. Participants were permitted to repeat the protocol provided that a minimum of one week had elapsed between exposures; however, repeat exposures were typically separated by several months when applicable. Participants underwent ultrasound imaging before, during, and after the hyperbaric exposure. In-dive imaging was performed using a research programmable ultrasound system during decompression, while post-dive venous gas emboli (VGE) were monitored using serial echocardiography examinations conducted every 20 min for at least two hours after surfacing using commercially available clinical ultrasound systems (see below).

### Hyperbaric exposure protocol

The hyperbaric exposure (dive profile) was chosen as an experimental profile previously characterized by the U.S. Navy Experimental Diving Unit (NEDU) that resulted in substantial post dive VGE but no DCS across 96 human dives (32). The pressure profile in the chamber consisted of a maximum depth of 132 feet seawater (FSW) or 505 kPa absolute (5.0 ATA) for a bottom time of 20 minutes (defined as the interval from start of pressurization to initiation of departure from maximum depth). The decompression phase was linear with a stop at 20 FSW (162 kPa) for 9 minutes during the return to surface (Fig. 1b). In contrast to the original NEDU profile, a brief stop at 30 FSW (193 kPa) for approximately 1 minute was included to transfer the subjects from the primary chamber to the adjacent ultrasound chamber. Rates of compression and decompression were approximately 50 FSW/min (153 kPa/min) and 30 FSW/min (92 kPa/min), respectively. Participants were instructed to alert the in-chamber tender if they had trouble equilibrating their ears during compression. Participants remained dry and breathed air throughout the hyperbaric exposure.

**Figure 1.**
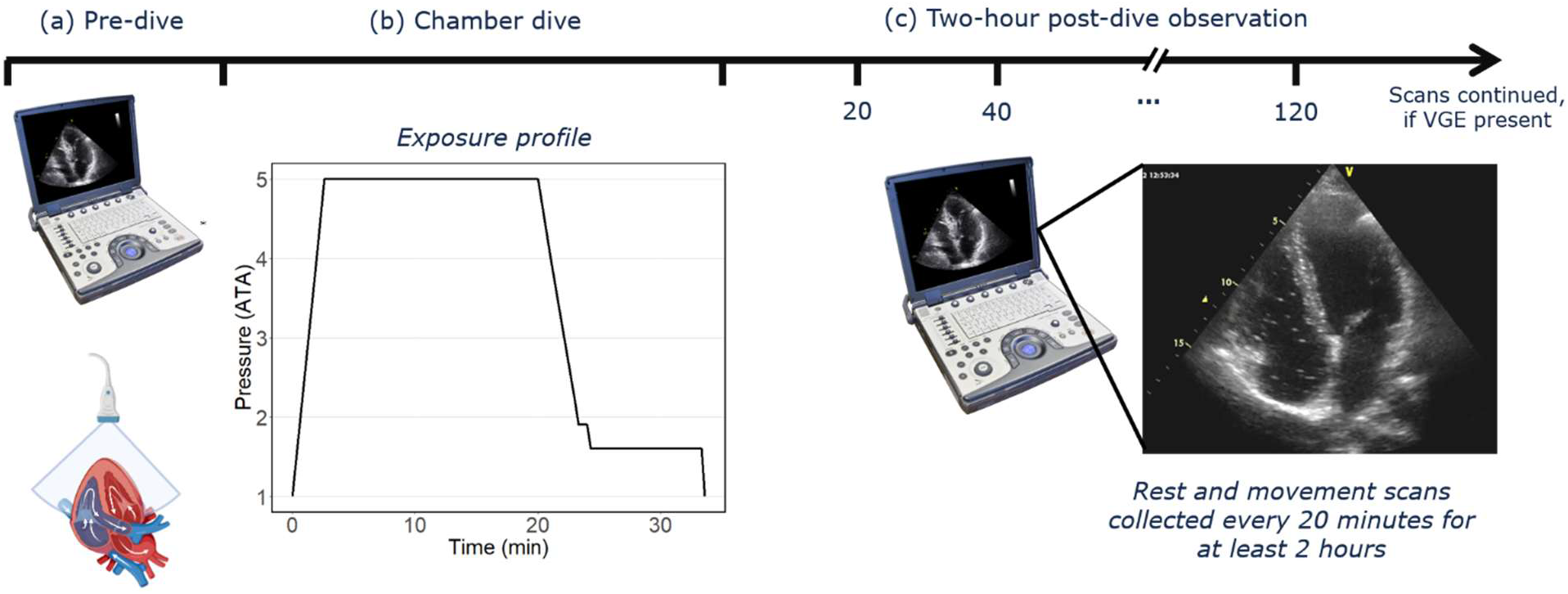
Experimental timeline for human dive trials. (a) Pre-dive baseline scans collected with clinical and research ultrasound systems. (b) Participants conduct dry hyperbaric chamber dive to 132 FSW for 20 minutes followed by staged decompression. (c) After exiting chamber, echocardiography is conducted for at least two hours to measure venous gas emboli (VGE) presence.

During the dive, participants underwent one of three protocol variations. Initially, participants remained seated at depth without exercise (***Non-exercising***, n = 9). After a mid-study review of VGE results demonstrated lower than anticipated peak post-dive VGE, participants performed light exercise on a cycle ergometer (***Exercising***, n = 20) during the bottom phase at 132 FSW, similar to the NEDU implementation (32). As part of a separate experiment, a brief pause of approximately 1-2 minutes at 20 FSW was added during the compression phase for some of the dives (***20-FSW pause***, n = 12). This pause was considered part of the bottom time and included within the 20-minute interval comprising time for compression and time spent at 132 FSW.

### Post-dive VGE monitoring and grading on echocardiography

After exiting the hyperbaric chamber, participants were monitored with ultrasound every 20 minutes for at least two hours (Fig. 1c), as recommended by current field consensus guidelines for VGE detection with ultrasound (17, 31). At each imaging time point, a trans-thoracic apical 4-chamber view was collected by trained research operators experienced in VGE image acquisitions with one of three commercially-available clinical ultrasound systems: the Mindray M8 Elite ultrasound system with an SP5-1s probe (Mindray, Mahwah, NJ), the Sonosite PX system (FUJIFILM, Bothell, WA) with a P5-1 probe, or the VividQ system with an M4S-RS probe (GE HealthCare, Chicago, IL). A recording of at least 10 cardiac cycles was collected in adult cardiac imaging modes for each participant both at rest and immediately following three forceful leg movements.

Echocardiography recordings were exported in audio video interleave (‘.avi’) format with frame per second (FPS) rates varying by system 25 FPS (Mindray), 45 FPS (VividQ), and 21 FPS (Sonosite). Videos were reviewed offline by a single, experienced VGE grader (JBC) at normal and half speed to ensure grading accuracy. Post-dive VGE were graded on the Eftedal-Brubakk (EB) 0-5 ordinal scale for both the rest and movement instances, ranging from EB grade 0 indicating no detectable bubbles to grade 5 corresponding to maximal bubble presence (Table 1) (31). The resting grade was assigned based on the sustained VGE presence across 10 consecutive cardiac cycles. The movement grade was assigned using the highest VGE presence maintained across six cycles occurring after initiation of leg movements. The shorter grading interval for movement grading has been described to better capture transient bursts of VGE that may not be sustained for 10 consecutive cardiac cycles (16). The peak post-dive EB grade for any instance (rest or movement) was identified for each participant, along with the time to peak grade and duration of detectable VGE presence post-dive.

**Table 1.** Eftedal-Brubakk (EB) Grading Scale.

| EB Grade | Definition |
| --- | --- |
| 0 | No bubbles |
| 1 | Occasional bubbles |
| 2 | At least 1 bubble every four cycles |
| 3 | At least one bubble per cycle |
| 4 | At least one bubble per cm<sup>2</sup> in all frames |
| 5 | “White-out”, individual bubbles cannot be discerned |

### Ultrasound system preparation for hyperbaric operation

The Verasonics V1 (Kirkland, WA, USA), a programmable research ultrasound system, enables custom real-time pulse sequence design and access to raw IQ channel data for receive signal analysis. For hyperbaric operation, the V1 system was placed into two custom-sealed plastic enclosures equipped with cable ports to allow wired communication between the host computer, V1 system, and peripherals (Fig. 2a). One container housed the host computer and the second housed the V1 system. All ports were covered with plastic lining and adhesive tape to prevent gas leakage through enclosure seams and cable interfaces (Fig. 2b). A connected monitor was entirely covered with a clear plastic bag to isolate it from the environment (Fig. 2b). System power was delivered by auxiliary 120-V outlets ported to the inside of the hyperbaric chamber. Early testing revealed that the V1 system was prone to rapid overheating when covered. To address this, pure nitrogen was added as a direct line and continuously supplied to each enclosure to aid system cooling and reduce the oxygen partial pressure within the isolated environment. Approximately 10 L/min of N_2_ gas was fed directly into each enclosure throughout operation in the chamber. The isolated environments were monitored for temperature and gas composition. A maximum oxygen level of 6% was established as the operational limit to minimize the risk of fire in the hyperbaric chamber environment, based on the National Fire Protection Association safety compliance codes (33, 34).

**Figure 2.**
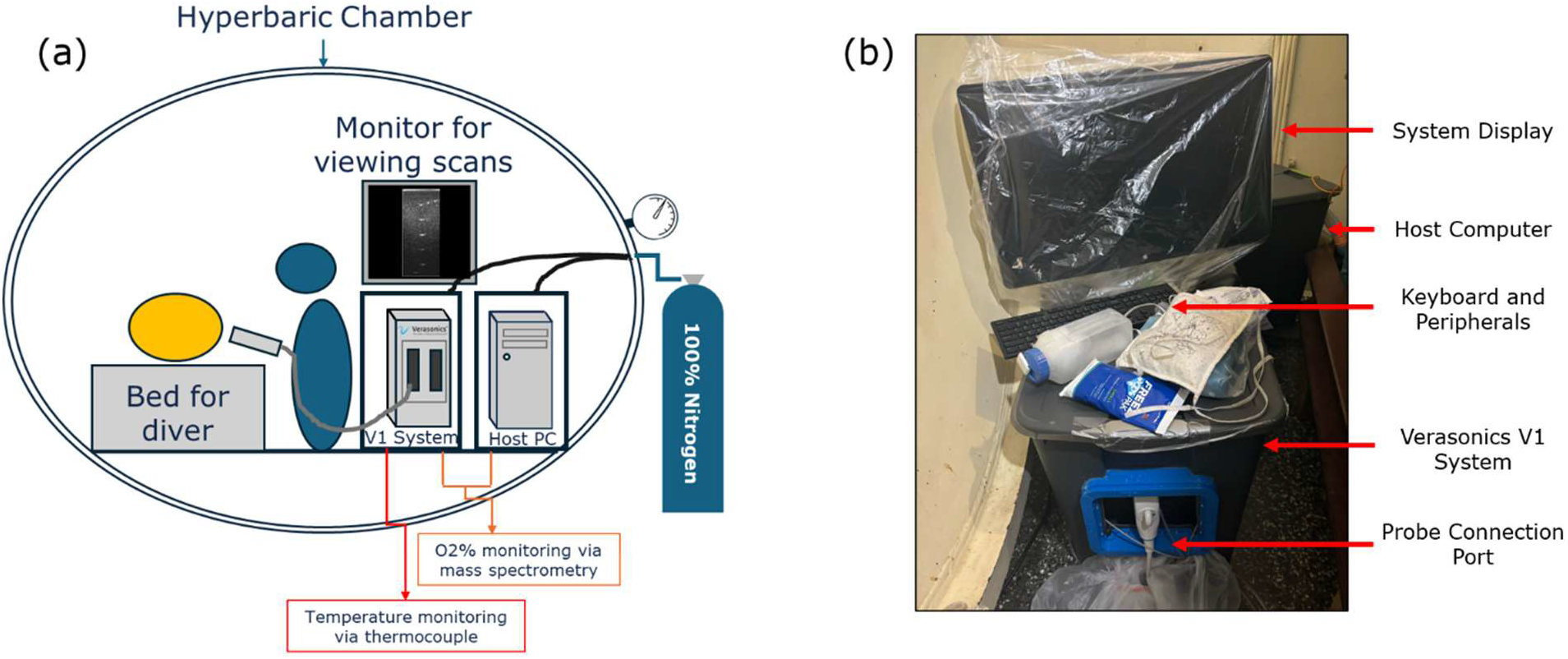
Preparation of research ultrasound system for hyperbaric environment. (a) Schematic of ultrasound station in the chamber with external nitrogen lines feeding the boxes. Temperature and gas composition are monitoring via thermocouple and mass spectrometry sample lines. (b) Verasonics V1 ultrasound system packaged prior to hyperbaric chamber dive with labels for different components.

### System testing at pressure

Ultrasound system operation and image acquisition performance were assessed at the surface and under hyperbaric conditions prior to use in human studies. System safety compliance was verified by monitoring temperature and oxygen partial pressure of the enclosure with a thermocouple and mass spectrometer (Perkin-Elmer, Shelton, CT), which sampled directly from the enclosures. System operation was evaluated by imaging a commercial ultrasound phantom (ATS, Bridgeport, CT) using the L12-3v probe at atmospheric pressure and at a simulated depth of 30 FSW (chamber pressurization to 193 kPa). A 3-angle plane wave imaging sequence spanning 20 degrees was used to a maximum imaging depth of 6 cm. Each acquisition consisted of a pulse inversion transmissions (PI+ and PI-) at 4 MHz followed by B-mode transmission at 8 MHz. B-mode and pulse inversion acquisitions were performed at a mechanical index (MI) of 0.2 and 0.3, respectively, corresponding to the maximum non-derated output values measured during hydrophone testing. Ultrasound data were collected in 3-s acquisitions at 30 frames per second (fps), totaling 90 frames per acquisition. The imaging protocol was repeated following chamber pressurization to approximately 193 kPa (30 FSW equivalent).

Ultrasound data acquired at the surface and at 30 FSW (192 kPa) were compared. Raw IQ channel data were beamformed offline using MATLAB (MathWorks, Natick, MA) and Verasonics’ delay-and-sum (DAS) beamformer (35). Beamformed ultrasound images were visually reviewed to assess consistency of image quality across pressure conditions. For quantitative analysis, regions of interest (ROIs) excluding hyperechoic phantom reflectors were identified on the beamformed images. The corresponding envelope-detected ultrasound data were extracted and averaged within each ROI for each cine acquisition using a custom MATLAB script. Mean ROI pixel brightness values obtained at the surface and at 30 FSW were compared to assess signal consistency under hyperbaric conditions.

### In-dive human ultrasound imaging

For a subset of the study group (12 dives performed by 10 individuals), ultrasound images were collected with the programmable ultrasound system pre dive and during the dive. When time allowed, post-dive scans were collected and the time relative to end of dive was noted. The Verasonics V1 system operated within the sealed enclosure system was used to capture baseline (predive at atmospheric pressure) and during-decompression (at 20 FSW) ultrasound recordings of participants. Gas composition in the two enclosures was monitored with mass spectrometry and continuous N_2_ gas was fed directly into each enclosure. Matched settings were maintained for each individual using the Verasonics with a 7-angle plane wave imaging sequence spanning 20 degrees was used to a maximum imaging depth of 6 cm. Each acquisition consisted of a pulse inversion transmissions (PI+ and PI-) at 4 MHz followed by B-mode transmission at 8 MHz. B-mode and pulse inversion acquisitions were performed at a mechanical index (MI) of 0.2 and 0.3, respectively. Ultrasound data were collected in 3-5s acquisitions at 30 frames per second (fps), totaling 90-150 frames per acquisition. For each imaging session, participants were either in supine position with one knee bent or fully prone, which was kept constant for each participant. Baseline scans were collected immediately before the dive and decompression scans were collected during the 9-minute decompression stop at 20 FSW. To evaluate consistency of probe placement and resulting ultrasound image data, a test-retest design was implemented. For nine participants, the pre-dive calf muscle scans were collected at least twice with the transducer fully removed and repositioned between acquisitions.

Ultrasound data collection during the dive was limited to the decompression phase beginning at the 192 kPa (30 FSW) stop. Operation of the packaged Verasonics V1 system during decompression required two operators, one to position the probe (VP) and the other to interface with the host computer (JBC). For in-dive imaging, the two ultrasound operators were pressurized to 30 FSW (192 kPa) in the ultrasound chamber, which shares a lock with the primary chamber, with the enclosed V1 system. After the primary chamber reached 30 FSW during decompression, the participants walked through the lock into the ultrasound chamber and immediately assumed the imaging position. Once the ultrasound chamber completed travel to 20 FSW (161 kPa), ultrasound scans were conducted on each participant. Recordings were acquired using the same protocol in both the pre-dive imaging session and during the dive: sagittal calf muscle, transverse calf muscle, and subclavian vein imaging when time permitted. Participants rotated through the imaging station until the end of the 9-minute decompression stop, occasionally allowing for multiple scans of each diver to be collected.

### Programmable ultrasound image analysis

Post-experiment, ultrasound IQ channel data were beamformed offline as previously described for phantom experiments and log-compressed to grayscale for visualization. Individual recordings were reviewed and trimmed for motion or buffer inconsistencies. Subclavian recordings were individually reviewed in slow motion for possible circulating VGE, which typically appear as discrete hyperechoic signals moving within the venous lumen.

Sagittal calf muscle recordings were selected for image brightness analysis. Recordings were trimmed to remove motion artifacts and ensure consistency across different imaging time points. The beamformed, envelope-detected ultrasound data were analyzed prior to log compression. B-mode and pulse inversion datasets were analyzed separately. An ROI was manually defined within the boundaries of the gastrocnemius muscle for each recording, and individual pixel values were extracted from each frame within the cine acquisition. For each frame, pixel values within the ROI were averaged to obtain a mean ROI intensity, and these values were subsequently averaged across all frames to obtain a single intensity value for the recording.

### Statistical analysis

After each dive, the peak post-dive EB grade was identified as the highest grade achieved at any time, regardless of type (rest or after leg movement). The median peak EB grade across all dives per exposure type (Non-exercising, Exercising, 20-FSW pause) were obtained and groups were compared for similarity using a Kruskal-Wallis test. The relationship between peak EB grade and time to peak per dive was described using Spearman’s rank correlation. For in-dive imaging, changes in mean ROI intensity between baseline and decompression scans were calculated separately for B-mode and PI datasets. The magnitude of brightness change from the decompression stop to pre-dive was calculated across the different scans for both B-mode and PI ultrasound data. The relationship between average brightness change and peak post-dive EB grades was evaluated using a Spearman rank correlation.

Probe placement test-retest reliability was estimated by acquiring multiple pre-dive baseline calf muscle scans in some participants (n=9). The standard deviation was obtained for each individual, then the relative percent variation was established as the ratio between standard deviation and average baseline value per participant. The average percent variation across all participants was defined as the error associated with probe positioning. This error was then propagated to differences between decompression and predive measurements, as well as percent changes, through standard error propagation algebra techniques (36).

## Results

### Dive profile and VGE presence

No DCS cases were reported across the three different dive exposure types totaling 42 human dives by 26 unique participants. One participant experienced middle ear equilibration difficulty and was removed from the chamber within 1 minute of pressurization. This participant was excluded from further measurement and analysis. Post-dive VGE trends varied across both exposure types and individuals (Fig. 3). Substantial inter-individual variability in VGE production was observed within each exposure group.

**Figure 3.**
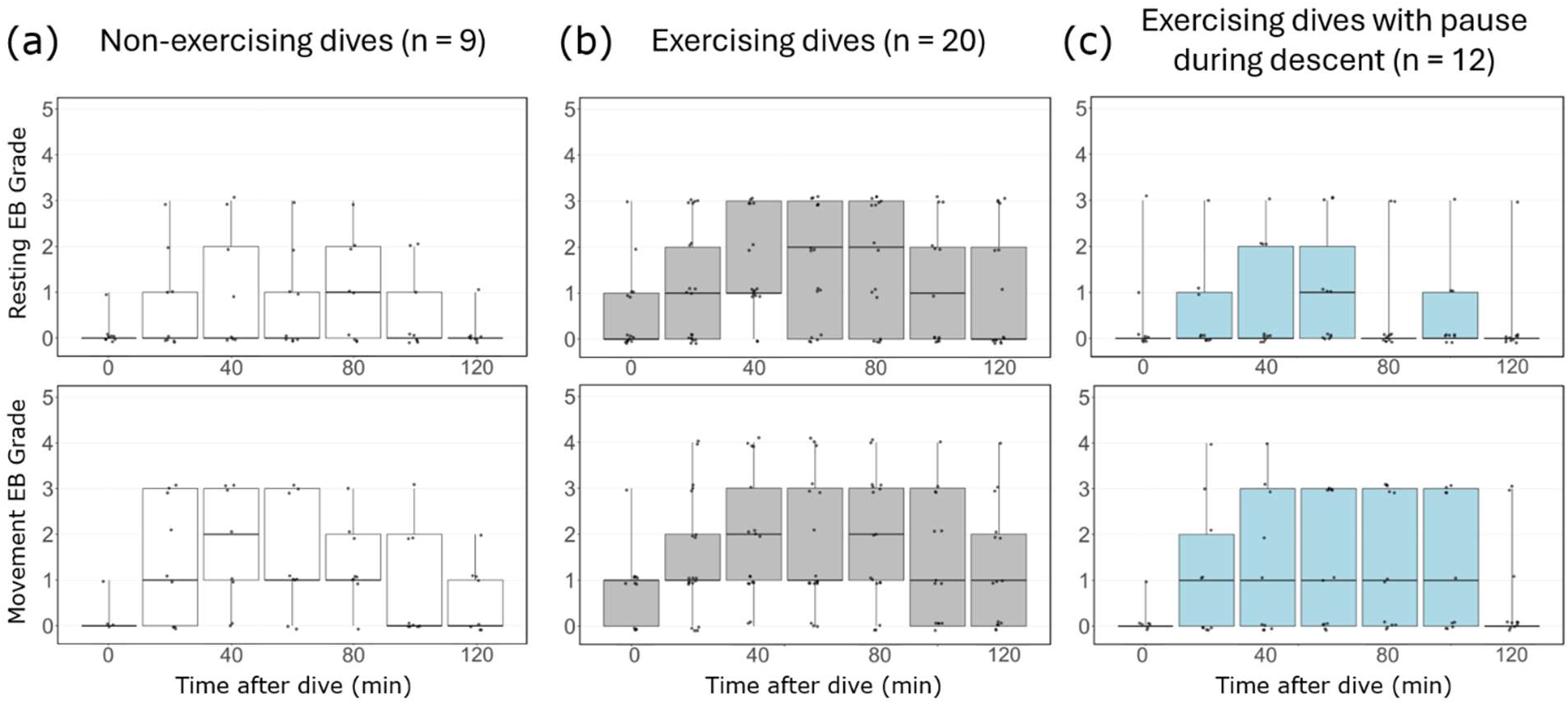
Post-dive VGE timelines for the varying exposures. The VGE grades at rest (top) and after movement (bottom) are included for non-exercising dives (a), exercising dives (b), and dives with a 20 FSW break during descent (c). Boxplots present the median and span the range for all measurements collected a given time post-dive.

Some participants during the exercising dives tended to exhibit greater EB grades and more prolonged VGE presence than non-exercising and 20-FSW pause dives (Fig. 3). The median peak post-dive EB grade was 2 for the non-exercising and 20-FSW-pause exposures, and 3 for the exercising exposure (Fig. 4a), with each exposure group found to have in a range of peak post-dive VGE spanning four EB grades. Comparison of groups found the difference was not significant with p-values (Kruskal-Wallis test) of 0.32 and 0.66 for EB grades and time to peak, respectively.

**Figure 4.**
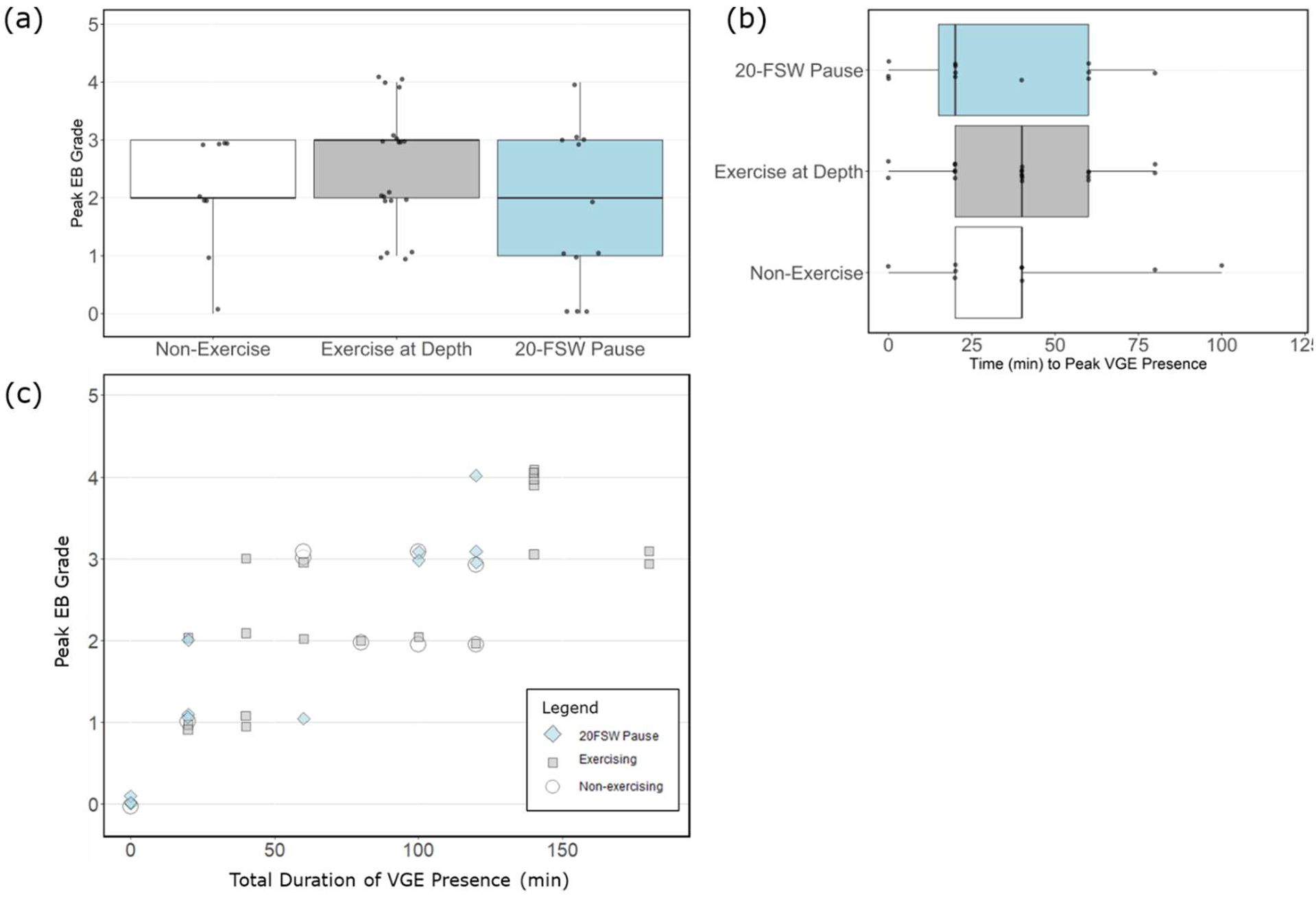
Summary EB Grades for the different dive exposures. (a) Peak post-dive EB grade for each dive. (b) Earliest time to occurrence for peak VGE presence for each dive. (c) Comparison between peak post-dive EB grade and total duration of VGE presence. All boxplots present the median and range for the data.

Peak VGE presence was typically found to occur within 1 hour of surfacing, with observed time to peak grade ranging from immediately post-dive to 2 hours after surfacing (Fig. 4b). Supplemental Information 1 provides a further breakdown between resting and post-movement scores for both peak grade and time of occurrence. Higher peak EB grades were associated with longer durations of detectable VGE (Spearman ρ = 0.82, P < 0.01, Fig. 4c).

### Ultrasound system operation at depth

After refinement of the isolated environment for the Verasonics V1 system, system operation was successfully maintained during tests at surface pressure (101 kPa) and the pressure equivalent of 30 FSW depth (193 kPa). Temperature and oxygen measurements were within acceptable ranges while continuously imaging a commercial ultrasound phantom for more than 30 minutes, exceeding the duration required for the planned decompression stop imaging period. Beamformed ultrasound images collected on the phantom during pressure testing are shown in Fig. 5a-b.

**Figure 5.**
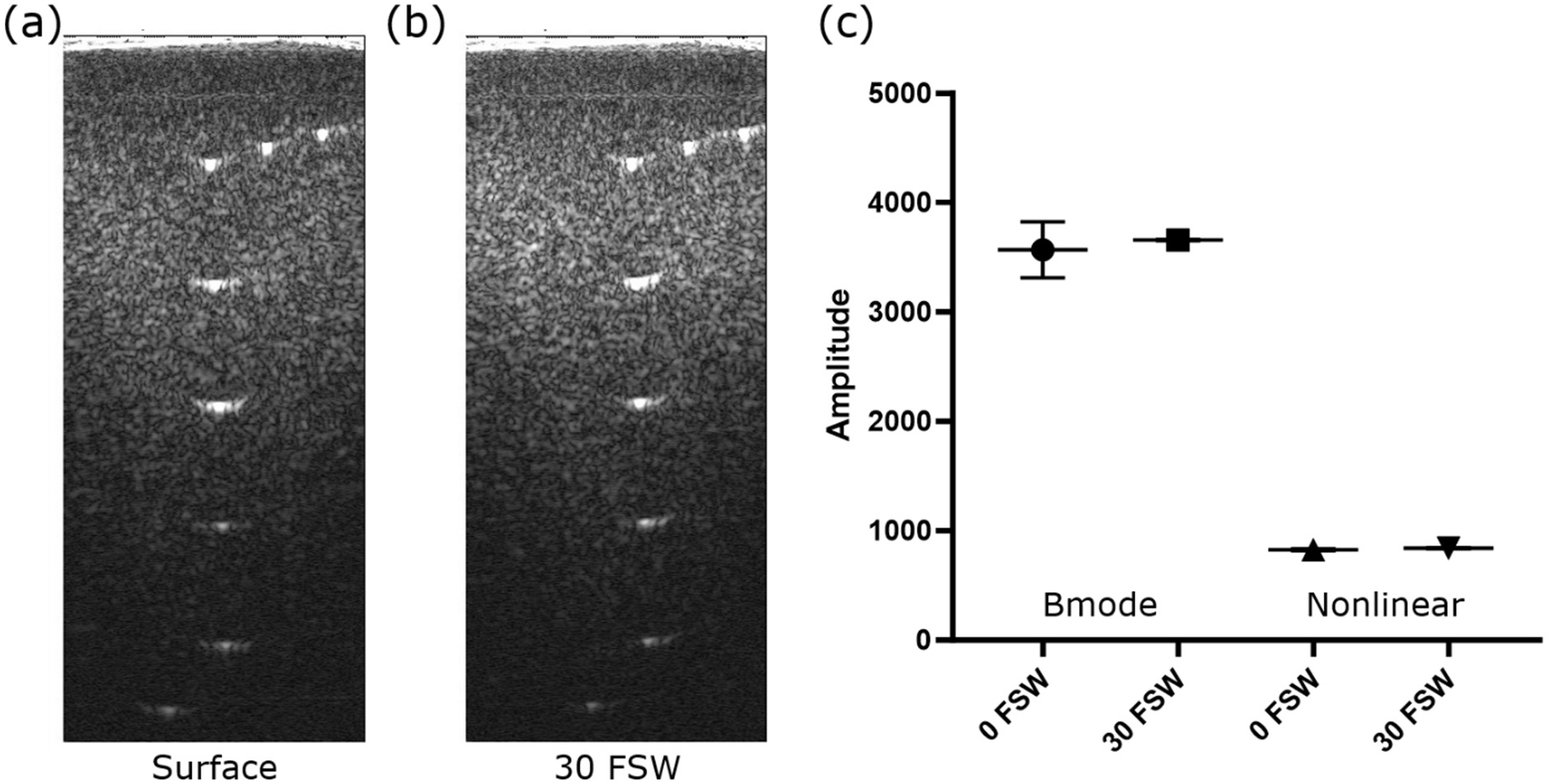
Ultrasound system phantom validation. Example phantom images collected at atmospheric pressure (a) and at nearly 2 atmospheres (b). (c) Brightness analysis of background from images collected in brightness mode and pulse inversion mode at the surface and depth.

Analysis of the envelope-detected ultrasound signal revealed a slight brightness change in the phantom ROIs in amplitude of 90 a.u. on B-mode and 13 a.u. on non-linear mode (Fig. 5c). These changes were small relative to the overall signal intensity and supported consistent system performance under hyperbaric conditions. Recordings of temperature and gas composition within the enclosure during the first trial with study participants are provided in Supplemental Information S2.

### Pilot human diver imaging during decompression

Ultrasound imaging was successfully performed before and during decompression in 12 hyperbaric chamber dives conducted by 10 participants. All participants but one were found to have VGE present on post-dive echocardiography within 2 hours of surfacing. Peak post-dive EB grades ranged from 0 to 4 for the 12 dives, with no reported DCS symptoms.

Imaging of the subclavian vein during decompression revealed circulating VGE in two participants (Fig. 6a-c). Both participants subsequently demonstrated VGE on post-dive echocardiographic examinations collected immediately after surfacing and reached a peak EB grade 3 during the post-dive monitoring period. A slowed recording of VGE in the left subclavian vein captured during decompression is included as Supplemental Information S3.

**Figure 6.**
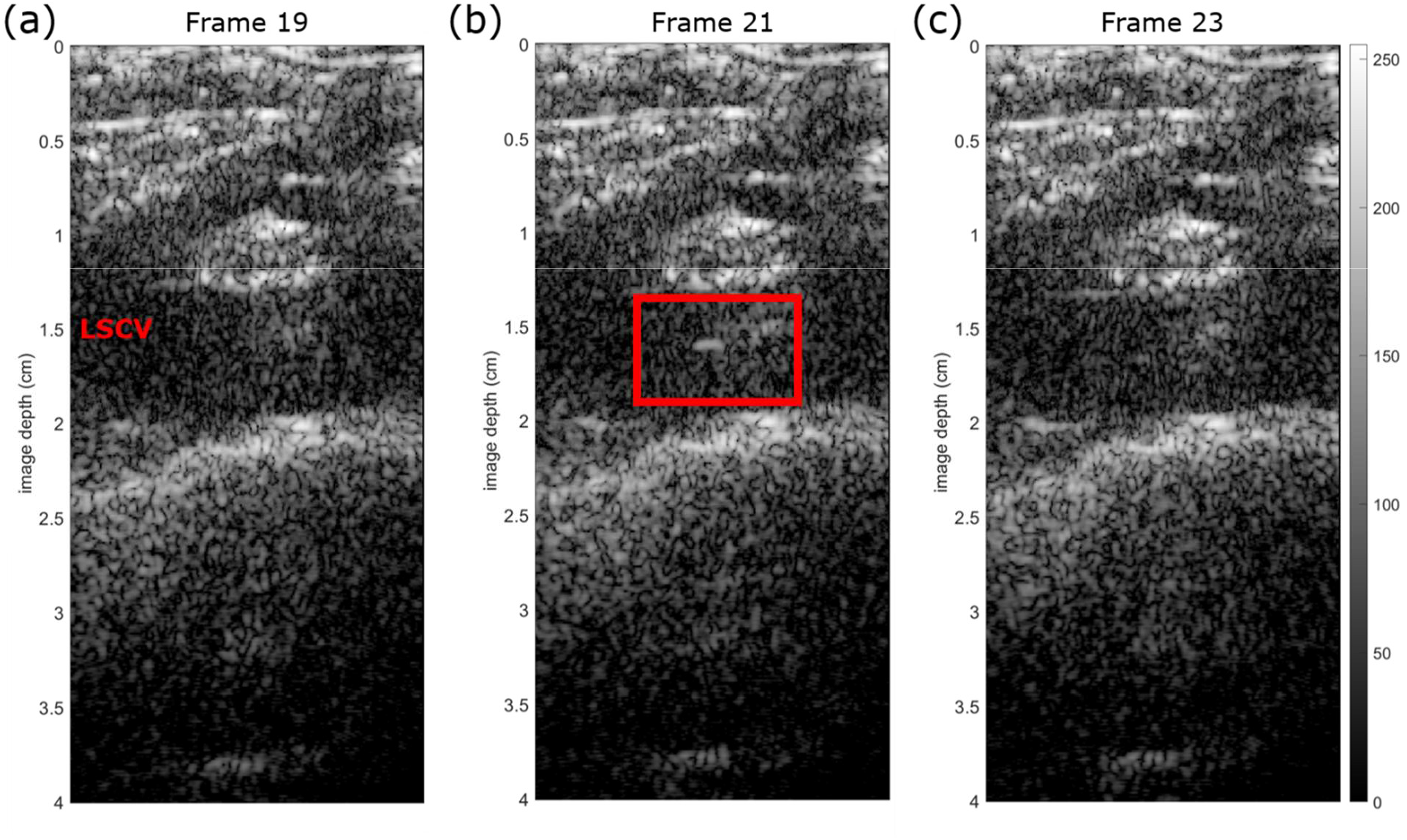
VGE detected during decompression in left subclavian vein (LSCV). (a) Frame of subclavian vein captured during decompression. (b) VGE subclavian vein highlighted by red box detected two frames after. (c) No VGE are visible two frames later.

### Calf muscle image brightness analysis

Probe positioning on the calf muscle parallel to the length of the fibula produced the largest set of consistent images for longitudinal comparison. Example calf muscle images from a participant pre-dive and during-decompression are presented in Fig. 7a-b. Ten of 12 dives demonstrated an increase in calf muscle signal intensity during decompression relative to baseline (Fig. 7c-d). A median peak post-dive EB grade 3 with no DCS cases for the 12 subjects with scans during decompression, with a range from 0-4. Signal intensity of in-dive calf muscle ultrasound data increased with an average 16.5% change for B-mode and 5.1% change for PI relative to baseline, while the phantom ROI had 2.5% B-mode and 1.6% PI changes. B-mode brightness change was not correlated with peak post-dive EB grade (Spearman ρ = 0.086, P = 0.79); however, a significant positive relationship was found when focusing on EB grades 0-3 (Spearman ρ = 0.64, P = 0.048). No correlation was found between PI signal change and EB grades (Spearman ρ = −0.45, P = 0.14), but a negative tendency was identified for EB grades 1-4 (Spearman ρ = −0.57, P = 0.066). The test-retest measurements on the subset of divers produced a relative uncertainty of 10.05% for B-mode and 3.55% for PI scans attributed to probe positioning during the pre-dive scans. Error propagation produced a 17.4% and 6.1% uncertainty for percent difference calculations of B-mode and PI scans, respectively. Across the 12 dives with during-decompression scans, an average signal increase of 1350 a.u. was found for the B-mode data and 48 a.u. for the pulse-inversion data.

**Figure 7.**
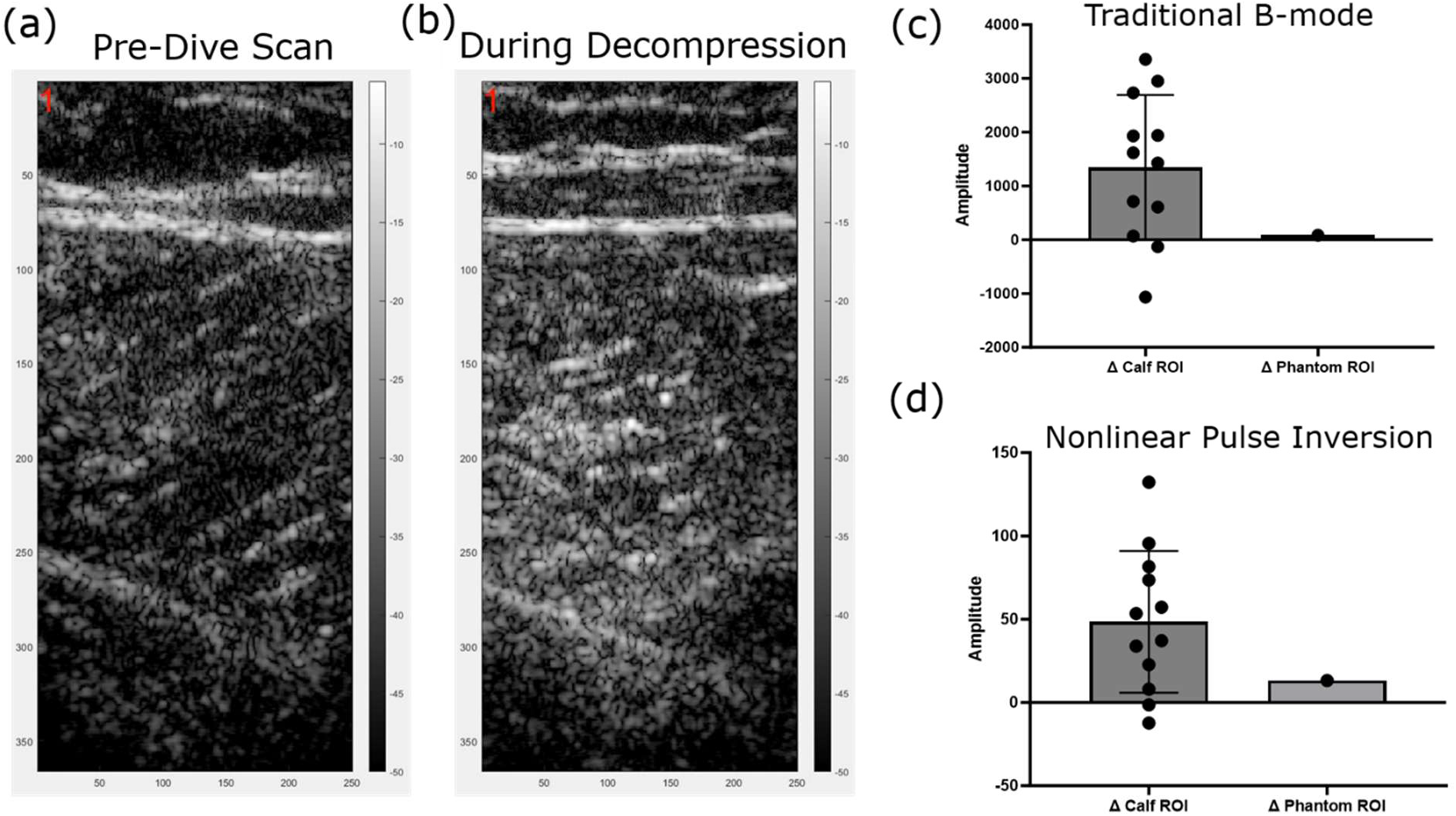
Calf muscle scans during decompression and brightness results. An example calf muscle scan from one diver including pre-dive (a) and during decompression (b). (c) B-mode and (d) Pulse Inversion image brightness analysis comparing the calf muscle scans collected from pre-dive to during decompression. The calf muscle images were typically brighter during decompression.

Two participants demonstrated decreases in B-mode ultrasound signal amplitude within the calf muscle ROI (magnitude 127 a.u. and 1060 a.u., Fig. 8). Divers with peak EB grade 3 were found to typically have an increased calf muscle brightness during decompression, relative to their pre-dive scan, but the two divers with peak EB 4 had a lower magnitude change (Fig. 8a). The PI mode produced a similar result to B-mode but in this case only participants with peak EB 4 showed a small decrease in amplitude during the dive (of lesser magnitude than the small increase in signal seen in the phantom data) (Fig. 8b). Calf muscle signal intensity increased from pre-dive to in-dive, then was found to decrease post-dive beyond the baseline value for all divers, except one (Fig. 9a-b). One individual, with peak EB 4 VGE, followed an opposite trend in calf muscle signal intensity with a decrease during the dive and increase two hours post-dive. Time, personnel and equipment limitations precluded consistent post-dive calf muscle measurements. Extended time-series plots per diver of B-mode and PI calf muscle scans can be found in Supplementary Information S4.

**Figure 8.**
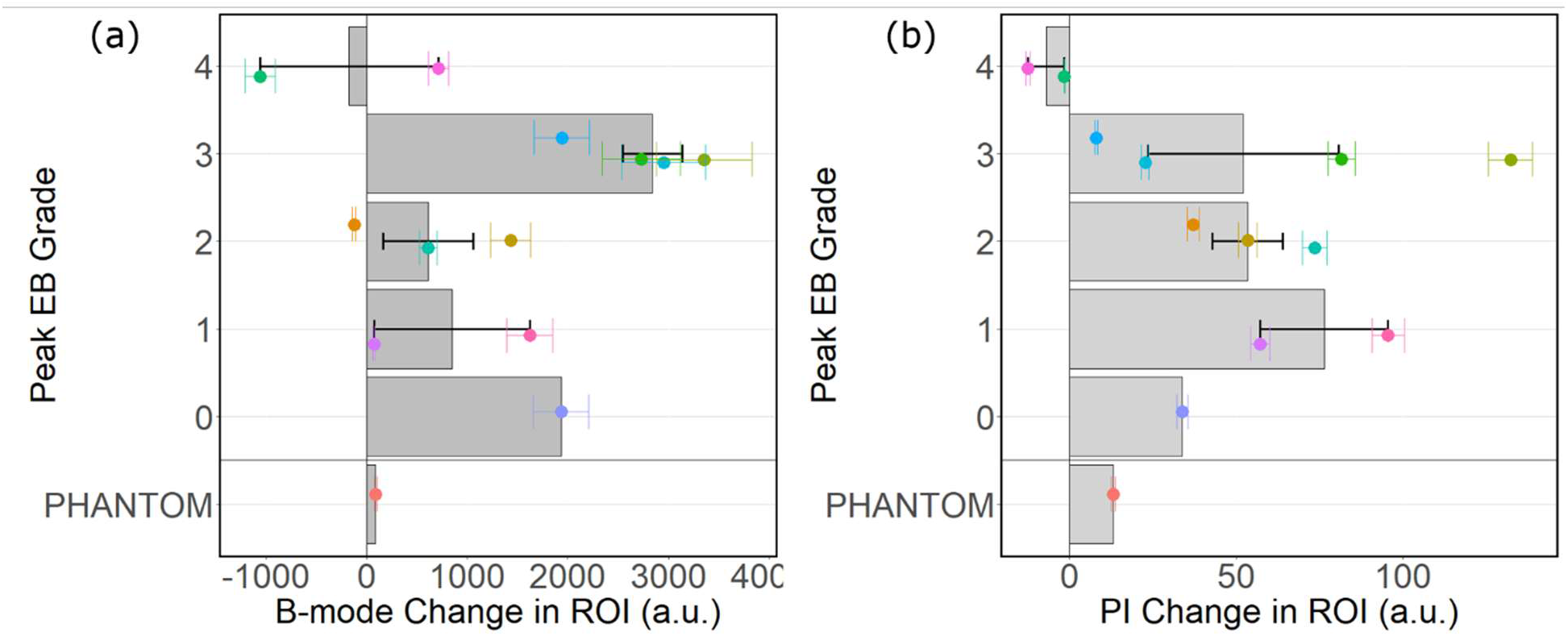
Comparison of calf muscle echo intensity change and peak post-dive EB grades. Each point represents a single diver for the peak EB grade against the change in calf muscle ultrasound signal from pre-dive to during decompression for (a) B-mode and (b) Non-linear mode. Change in ultrasound signal was measured on the envelope-detected, uncompressed data. The within subject standard deviation (σ_WS) comes from the test-retest images collected on six of the participants.

**Figure 9.**
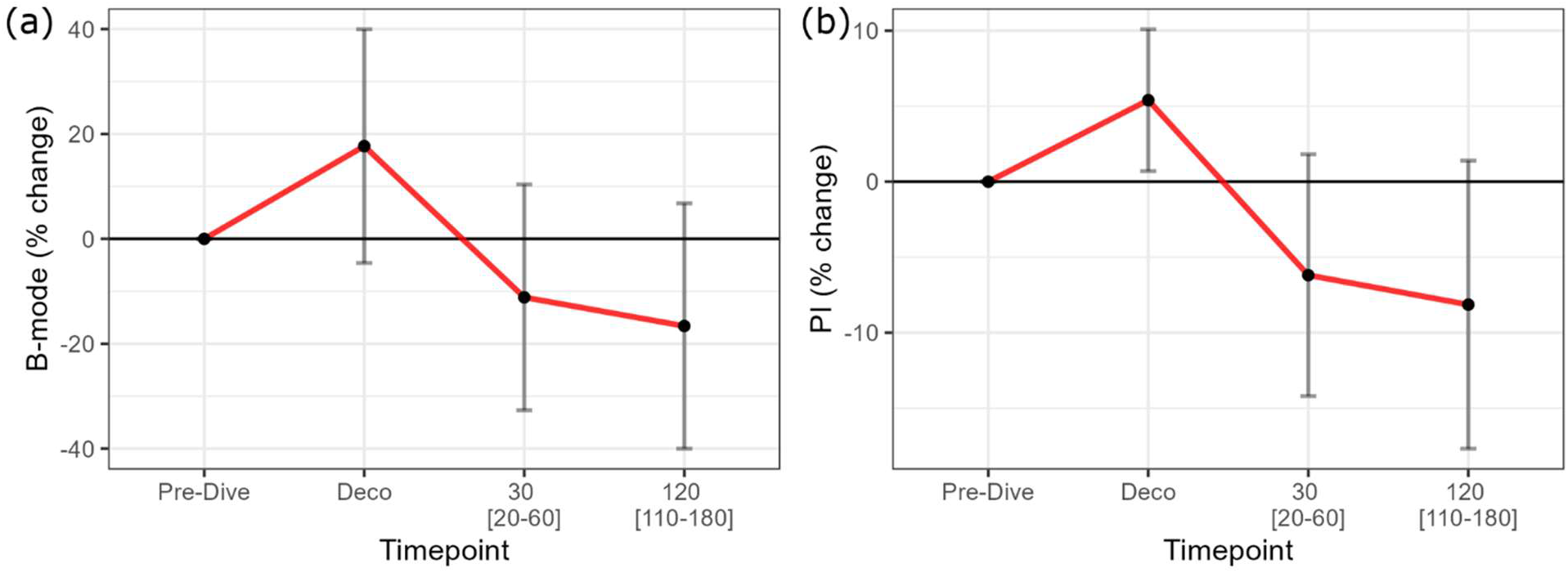
Average percent change of calf muscle brightness per timepoint for (a) Brightness mode (B-mode) and (b) Pulse Inversion (PI). Timepoints are ordered sequentially including pre-dive, during decompression, 30 minutes post-dive, and 120 minutes post-dive (brackets indicate time ranges for scans collected). Percent change is relative to pre-dive baseline scan and averaged across up to 12 participants at each timepoint.

## Discussion

Historically, ultrasound and physiological data have been accessible during a dive or hyperbaric exposure have been limited. After refinement of the enclosure system, the Verasonics programmable ultrasound system operated safely in the chamber. Reducing the risk of operating an electronic instrument in the setting of increased oxygen partial pressure was challenging to overcome as it required isolation from the environment. Preliminary testing revealed that the system was prone to overheating inside the container, which was remedied by implementation of a constant flow of 100% nitrogen gas, which greatly extended the available imaging time. The workflow we developed for operation of a programmable ultrasound system during hyperbaric exposure allows for collection of the raw, unprocessed receive signal for post-study reconstruction and analysis with maximum flexibility. Previous studies have identified image brightness trends for pre- to post-dive using clinical systems in animals and humans (27, 28). However, ultrasound system manufacturers regularly implement automated image adjustment using a proprietary algorithm to produce an optimal image for clinical review, which may arbitrarily adjust the image brightness. This problem does not impact our programmable system as the uncompressed, unprocessed receive signal can be analyzed. Preliminary testing found that the time to save a recording was longer in the box at pressure than out of the box at the surface, which we attribute to the warming of the isolated environment as the system continued to run. To counter this and reduce time to save each scan, a smaller number of plane wave angles were collected for compounding and shorter frame buffers.

This study represents the first use of a programmable ultrasound system to acquire and quantitatively analyze ultrasound data during decompression in human participants. We were successfully able to detect VGE with two-dimensional ultrasound imaging in the subclavian vein during the decompression phase. Both individuals with VGE in-dive were found to have VGE on echocardiography immediately after completion of the dive and reached an EB grade 3 during observation, indicating elevated DCS risk. Due to time constraints, we were unable to collect ultrasound data of the subclavian vein in the EB 4 divers, so we cannot determine if VGE presence occurred during decompression for those participants. However, we noted that both peak EB 4 divers presented with VGE on the echocardiography scan collected immediately after the divers exited the chamber. Peak VGE presence typically did not occur for at least 20 minutes after the dive, so it is not expected that the maximal EB grade would have been identified during-the-dive. The detection of VGE in-dive affirms that decompression physiology is dynamic and reinforces the limitation of traditional post-exposure measurements.

This study also represents the first use of a programmable ultrasound system in a hyperbaric environment, providing access to information-rich, pre-processed signal. Ultrasound recordings captured of the standard commercial imaging phantom demonstrated consistent behavior of the Verasonics at the surface and at increased chamber pressure. The calf muscle ultrasound measurements revealed a consistent increase from pre-dive to in-dive for both B-mode and PI, a novel implementation and finding for the field. Preliminarily, the relationships between B-mode and PI with peak EB grades indicate the calf muscle measurements in-dive may have predictive power for the DCS risk of the individual, based on the established surrogate endpoint (16, 18). The conflicting trends for B-mode and PI to peak EB grades may be attributed to the varied imaging techniques, where PI isolates non-linear (microbubbles) from linear (such as tissue or larger bubbles) (31, 37). In our case, it is possible that divers with more post-dive VGE had early bubble growth, producing brighter B-mode images than PI images due to signal attenuation. The two divers with peak EB grade 4’s were found to a slight decrease in calf muscle ultrasound signal in-dive. While unexpected, this trend is similar to two preclinical studies that measured ultrasound image brightness in tissues and found that those with severe DCS tended to have darker images post-dive (28, 29). The image darkening was attributed to acoustic shadowing due to large gas presence (28, 29). It is possible that we observed a similar mechanism in the calf muscle scans, since a peak EB grade 4 corresponds with substantially higher VGE loads. The calf muscle imaging technique may lay the foundation for a future wearable ultrasound device for in-dive physiological monitoring, with correlation to a known post-dive endpoint.

Additional post-dive calf muscle measurements were collected post dive, allowing for time-series tracking. The consistent trend of increased calf muscle signal in-dive followed by decrease post-dive, regardless of post-dive VGE presence, supports the premise of a physiologically driven phenomenon being detected. Speculatively, it is possible that bubble formation and growth occurred during the dive, causing the ultrasound signal increase, followed by bubble transport and removal during the post-dive phase, triggering a signal decrease. The exact mechanism of VGE formation is yet to be elucidated, but a prevailing theory is the presence of bubble ‘micronuclei’ that act as stable growth sites for VGE development prior to release in the vascular system (38). Interestingly, the participant with an opposite trend of a calf muscle signal change (decrease in-dive, increase post-dive) also had a high VGE presence (EB4), which may be caused by differences in bubble formation and timing for greater inert gas presence.

The increased ultrasound signal during-the-dive indicates a heightened echo intensity of the calf muscle, which could be attributable to various mechanisms. Previously, Swan *et al*. detected a post-dive change in non-linear ultrasound signal in the muscle of a porcine model and theorized it may be related to decompression bubble formation (30), suggesting the change in ultrasound signal of the calf muscle may represent early bubble growth in our data. A pilot human study in seven participants also previously showed an increase in cardiac nonlinear ultrasound signal post dive followed by a decrease below baseline at 120min (39). An alternative explanation to the ultrasound signal change that we measured could be tissue inflammation instigated by exercise at depth and the hyperbaric exposure, both of which have been linked to inflammatory pathways (40, 41). A review on echo intensity within skeletal muscle measured using B-mode ultrasound found studies measuring correlation with muscle quality, glycogen levels, and inflammatory edema within the target region, albeit with some degree of uncertainty (40). However, exercise induced muscle echo intensity changes start multiple days after maximal eccentric arm actions (42), suggesting that the signal change detected in our study is not likely to be due to exercise response alone. ultrasound data was acquired using manual probe positioning. While the same operator conducted all scans for all participants and targeted individually consistent anatomical landmarks, some variability was unavoidable due to small differences in probe pressure, orientation and tilt (43, 44). We accounted for this by implementing a test-retest process for the pre-dive scans on half the subjects, allowing for quantification of deviation that is likely due to probe placement. May *et al*. have previously demonstrated intra-operator consistency when collecting ultrasound images of muscles for fascicle length measurements using a test-retest methodology but suggested standardizing the location with strict positioning protocols (45). Future work will include method refinement and quantitative image similarity assessment to ensure the same imaging plane is compared for the different scans.

Our replication of the dive profile to 132 FSW for 20 minutes of bottom time produced similar post-dive VGE results to the U.S. Navy Experimental Diving Unit (NEDU) (32), with no cases of DCS observed in either study. The exercise exposure yielded a median peak EB grade of 3, matching the median value reported from the 96 NEDU dives in Andrew & Doolette (32). However, the prior study found a median EB grade 3 at all post-dive measurement times (32), which was not the case in our data. Several factors may have contributed to these differences, including limited control of exercise intensity among participants and the use of dry chamber exposures rather than the immersed dives performed by Andrew & Doolette (32). Separately, others have shown that conducting a similar profile in a dry hyperbaric chamber produces less VGE than the same profile in wet field conditions (46), another possible factor for inter-study differences.

The non-exercising profile was initially selected for the study but produced lower peak VGE grades than the NEDU results, prompting the addition of light exercise during the bottom phase. Previous studies have demonstrated an impact of exercise during a dive on post-dive VGE (47–49), which may have contributed to differences in total amounts of inert gas up-taken on while at depth and hence the variances of post-dive VGE within our study. The contribution of varying exertion levels could not be discerned as the VO_2,max_ for each individual and cycle ergometer wattage throughout the dive were not measured. Considering the variability in peak VGE presence for the non-exercising dives, it is likely that exercise is not the sole factor in VGE outcome. The 20-FSW pause in decompression was implemented later in the study and was found to cause some participants to have a lower VGE peak and shortened post-dive VGE presence. The 20-FSW pause was counted as bottom time, thereby reducing the total time the divers spent at 132 FSW, which we attribute to being the main factor in lowering VGE presence. While a significant difference was not found for peak EB grades between the three exposure conditions, the data suggested exposure-dependent differences that may not have been fully resolved because of limited sample size. Doolette *et al.* previously estimated that a sample of 50 dives per arm would be necessary to exceed 80% power for a single VGE grade level difference (50), a threshold that we did not reach across the entire pilot study and a caveat to the VGE outcome interpretation.

Interestingly, we found that individuals with higher peak EB grades also exhibited a longer duration of VGE presence post dive. Previous work has discussed the correlation between peak EB grades and DCS risk at a population level; however, researchers typically stopped measuring VGE after identifying that VGE presence had declined from the peak (10, 51), and would be unable to estimate the total bubbling duration. The correlation between peak EB grades and total duration of VGE lends credence to VGE as a surrogate marker of decompression stress for a given dive exposure. Additionally, this positive relationship may help to explain the known variability in post-dive VGE grades (10, 52, 53). Under the base assumption that total ‘dose’ of inert gas taken on during the dive is the same, then one might propose a negative relationship between peak VGE presence and duration since the same volume of inert gas would need to exit the body. However, our results suggest the opposite relationship, indicating that the total ‘dose’ of inert gas taken on during each dive differed, causing a different post-dive VGE trend. Overlap in peak EB grades and VGE durations was seen for the different exposures (non-exercising, exercising, 20-FSW-pause), demonstrating that the modifications did not fully explain individual differences. Additional studies to better differentiate intra- and intersubject variability in VGE, and their relationship to variations in dive profiles, are warranted.

VGE presence remains the only marker related to DCS outcome over thousands of human dives (16, 18); however, echocardiography presents practical limitations outside of the research lab, most notably the required proficiency for probe positioning (31). The approach of ultrasound signal analysis of different tissue regions during-the-dive in the decompression research field may offer a complementary path towards estimating DCS risk with the potential for future automation and passive monitoring. The placement of the probe in the calf region with an automatable echo intensity analysis method could offer an easier implementation than the standard echocardiography and could be compatible with a wearable format (54). Undersea and space environments are remote and latency in data transfer could inhibit the analysis of echocardiography by a trained expert, further supporting the need for onboard, real-time feedback for the user. Operational divers and astronauts are regularly burdened with materials and tools necessary for the task at hand, thereby requiring any additional equipment to be strictly necessary for life-support. These findings warrant further data collection and advanced ultrasound signal analysis techniques to expand this pilot testing.

## Conclusion

Our results demonstrate for the first time the feasibility of using a programmable ultrasound system to safely acquire and quantitatively analyze ultrasound data during decompression in human participants inside a hyperbaric chamber. In addition to enabling in-dive physiological measurements, the workflow developed here provides a practical framework for operation of programmable ultrasound systems under hyperbaric conditions, potentially supporting future investigations in other hyperbaric medicine applications. Replication of a previously characterized NEDU dive profile produced post-dive VGE trends consistent with prior reports while demonstrating modest modifications to the exposure profile altered post-dive VGE responses. These findings further highlight the substantial inter-individual variability in decompression outcomes, even among participants undergoing similar exposures. VGE were detected in the subclavian vein during decompression in two participants using the programmable ultrasound system, both of whom also demonstrated VGE immediately upon surfacing as assessed by conventional apical four-chamber echocardiography. To our knowledge, this represents the first report of VGE visualized in the subclavian vein using two-dimensional ultrasound imaging during human decompression.

Most participants demonstrated increased calf muscle ultrasound brightness during decompression relative to pre-dive measurements. Signal decreases were not observed beyond the variability measured in phantom and test-retest analyses, suggesting that the observed brightness changes were unlikely to be explained solely by system drift or probe repositioning. Although the physiological basis of this signal change remains uncertain, these findings indicate that localized tissue ultrasound measurements during decompression capture a reproducible response that warrants further study. Future work will focus on refining imaging methodologies, expanding data collection to additional anatomical regions, and evaluating candidate measurement sites for future automated or wearable systems designed to monitor decompression stress in operational environments.

## Supporting information

Manuscript

## Notes

### Competing Interest Statement

J.B.C., O.O., F.Y.Y and V.P. are inventors on a provisional patent related to real-time decompression stress control with ultrasound. Additionally, O.O. and F.Y.Y employed by and/or have ownership in Clearsens, Inc, a company that has licensed CMUT technology they are co-inventors in and is commercializing wearable ultrasound for different applications. Other authors declare no conflict of interest.

### Funding

- Office of Naval Research: N000142012590, N000142312548
- United States Department of the Navy: N21A-T013-0040 phase II grant

