## Supplementary material for "Human decompression in real time: programmable ultrasound imaging during hyperbaric exposure": Manuscript

### Supplementary Information

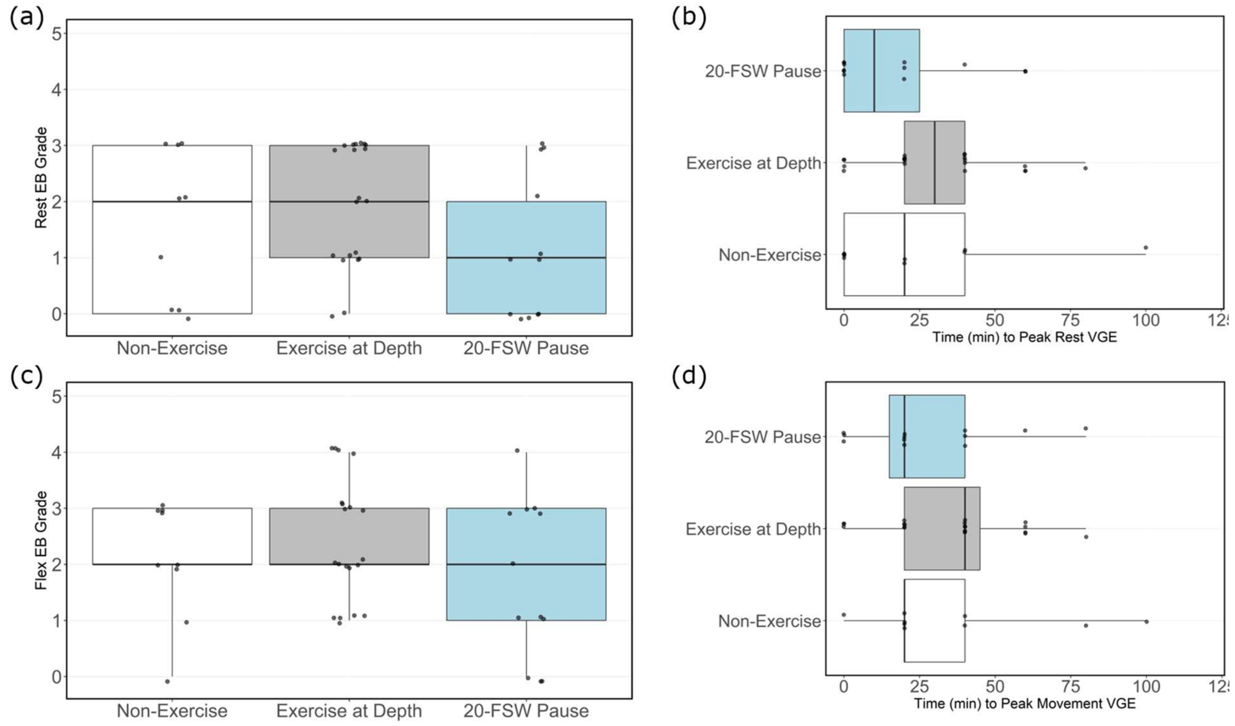


S1. Peak post-dive EB grades for rest and flex separated. The peak resting grades (a) and time of occurrence (b) include a single point for each dive exposure. Movement peak grades (c) and time of occurrence (d) are also included. All plots are separated by dive exposure.

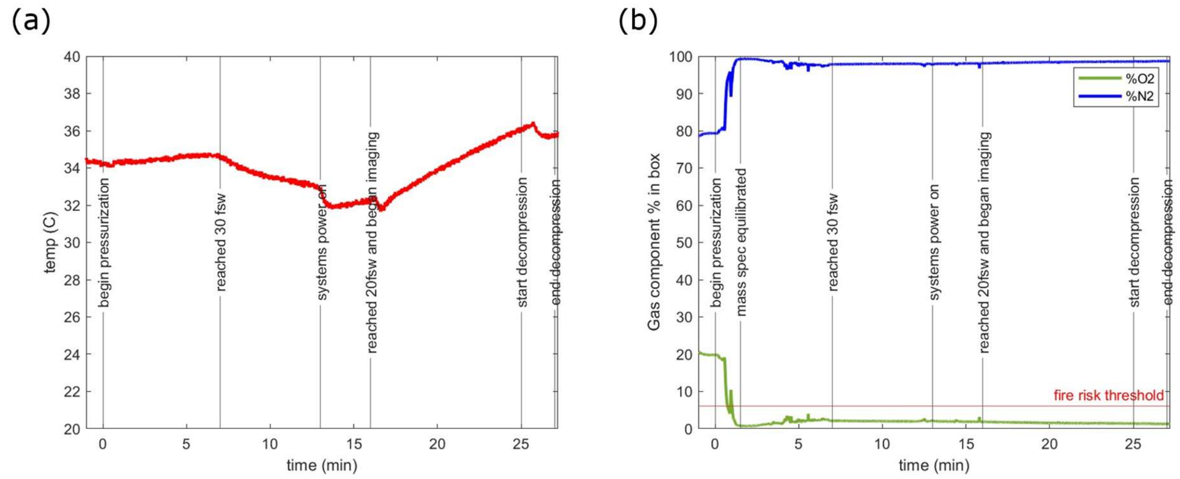


S2. Verasonics V1 box measurements during first dive with participants. Plots show the evolution of temperature (a) and gas composition (b) from the start of pressurization to the end of decompression with annotations. Notes on both plots indicate relative time points for the dive exposure and ultrasound system operation. The mass spectrometry system has a lag from the start of pressurization until accurate readings are obtained.

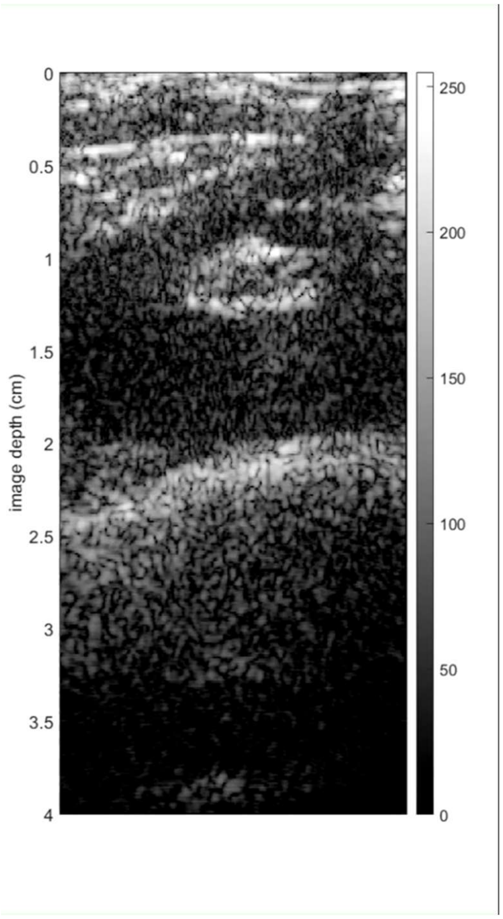


S3. Video captured during decompression of VGE in left subclavian vein. Recording originally captured at 30 frames per second but has been slowed to 10 frames per second.

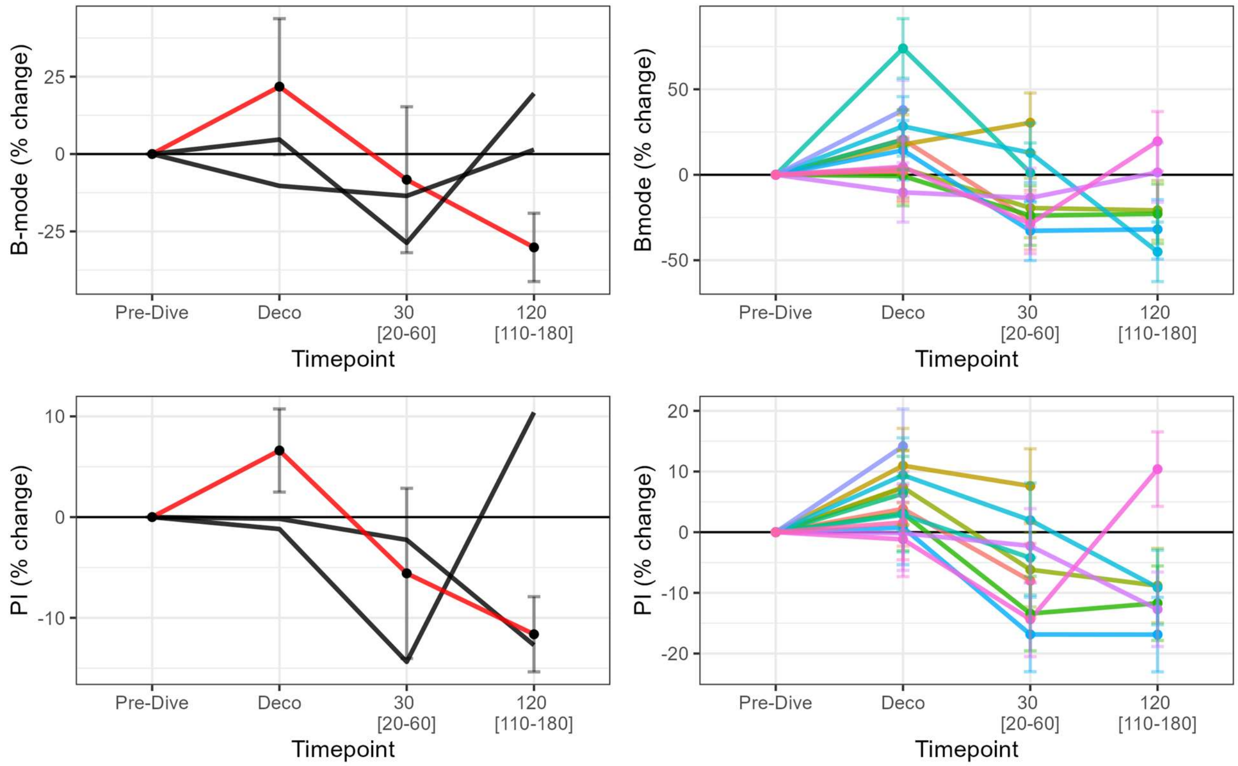


S4. Calf muscle signal intensity time series plots for B-mode (top) and Pulse Inversion (bottom) imaging schemes. Values represent the envelope-detected beamformed ultrasound data. Measurements from left to right are: pre-dive, during decompression, post-dive 1 (median 30 minutes, range 20-60), and post-dive 2 (median 120 minutes, range 110-180).
